# Planarians Develop Radiotolerance to Recurrent Ionizing Radiation Exposure

**DOI:** 10.1101/2024.10.25.620371

**Authors:** Paul G. Barghouth, Benjamin Ziman, Eli Isael Maciel, Peter Karabinis, Salvador Rojas, Natasha M. Flores, Edelweiss Pfister, Néstor J. Oviedo

## Abstract

Exposure to ionizing radiation can induce DNA fragmentation, leading to double-strand breaks, the most toxic form of DNA damage. Some organisms have developed mechanisms to overcome the adverse effects of ionizing radiation by enhancing DNA protection and repair. However, the underlying mechanisms driving radiation resistance to maintain genomic integrity and function remain poorly understood. Here, we provide evidence for the development of radiotolerance in the flatworm planarian *Schmidtea mediterranea*. We implemented a strategy to select animals capable of overcoming repeated rounds of ionizing radiation exposure. We demonstrate that planarians initially exposed to higher amounts of sub-lethal radiation can adapt, gaining the ability to recover reproductive capabilities faster than animals initially exposed to lower amounts of sub-lethal radiation. Our findings show that DNA integrity is reestablished in about one week after five cycles of sub-lethal ionizing radiation exposure. Planarian stem cells, known as neoblasts, can repair repeated DNA double-strand breaks by activating Rad51-mediated homologous recombination. The expression of the neoblast marker *smedpiwi-1* and the mitotic activity reach levels similar to unirradiated animals between two and three weeks post-radiation. We describe that planarians develop radiotolerance through recurrent ionizing radiation exposure over several years and survive without apparent functional or morphological defects for an undetermined time.

## 1. INTRODUCTION

Exposure to ionizing radiation (IR) can cause damage to cells and tissues[1]. IR has sufficient energy to displace electrons and modify the structure and function of molecules. For example, IR is known to alter DNA structure and cause fragmentation in single (SSB)- and double-stranded breaks (DSBs)[2-4]. DSBs are the most toxic forms of DNA damage, and cells have dedicated surveillance mechanisms to detect and repair the damage[5-7]. Indeed, evolution has imprinted an array of cellular mechanisms to recognize and aid in the repair of damaged DNA, such as cell cycle checkpoints, base excision repair for replacing a few nucleotides on a single strand of DNA, or non-homologous end joining (NHEJ)/ homologous recombination (HR) for the repair of SSBs and DSBs, respectively[7, 8]. If the IR-induced structural damage is extensive, cellular death decisions are implemented to prevent DNA mutations and cancer. These physical principles of IR are widely exploited in the clinic as radiotherapy to treat cancers, industry, research, and agriculture[9].

Despite the harmful effects of IR, some organisms have developed ways to adapt and survive in its presence. Indeed, radioresistance (RR) is widely found among organisms in three domains of life[10-12]. *Deinococcus radiodurans* and *Rubrobacter radiotolerans* display high Me/Fe ratios, which provide RR, allowing them to withstand 20 times more radiation than *Escherichia coli[13-15]*. In addition, *D. radiodurans* show enhanced HR DNA damage repair[16]. The fungus *Ustilago maydis* also includes activated forms of HR to develop RR[17]. The basidiomycetous fungus *Cryptococcus neoformans* has survived harsh radiation conditions, such as those at the Chernobyl nuclear power plant[18, 19]. Tardigrades, one of the smallest invertebrate organisms, are known for surviving high doses of radiation and chemical insults[12, 20, 21]. To identify the molecular mechanisms involved in radiation resistance, transcriptomic studies on tardigrades have identified overexpressed genes in response to IR exposure[20, 21]. Although the mechanisms driving RR are unclear, the evidence suggests that DNA protection and repair mechanisms may combine to maintain DNA integrity in tardigrades[20-22]. These findings suggest that RR could be part of a selection process that builds over time. We aim to introduce an experimental strategy to address how evolutionarily preserved DNA repair mechanisms may contribute to radiation resistance in adult tissues.

The planarian *Schmidtea mediterranea* is a flatworm with abundant pluripotent stem cells known as neoblasts. The neoblast is the only cell capable of dividing in the body and can differentiate into more than 100 possible fates, including muscle, neural, epithelial, etc.[23-27]. Thus, neoblasts are essential to support tissue regeneration and the continuous cellular turnover that can extend the organism’s lifespan for many years[28-30]. The asexual *S. mediterranea* strain used in this study reproduces by fissioning when physiological (e.g., animal size), environmental (e.g., number of animals, temperature), and metabolic (e.g., nutrient availability) cues are optimal[29]. Colony expansion occurs when a single planarian undergoes fission, regenerating missing parts in about one week and creating clonal organisms. Under optimal laboratory culture conditions, the fissioning events may take weeks to repeat on the same animal. Thus, *S. mediterranea* provides relatively rapid growth of clonal lines fueled by adult stem cells.

Experimental results from the last century demonstrated that IR exposure could eliminate the planarian neoblasts[31]. More recently, it has been established that a lethal IR dose (i.e., above 20-30 Gy) permanently eliminates neoblasts, leading to animal death in about three weeks[32, 33]. On the other hand, exposing planarians to IR less than 17.5 Gy is considered a sub-lethal dose that transiently eliminates neoblasts for about 24 hours, followed by a gradual neoblast repopulation[23, 34-37]. The molecular mechanisms driving neoblast repopulation after sub-lethal IR are not well understood. However, the extraordinary evolutionary conservation of DNA repair mechanisms in *S. mediterranea* mediates neoblast repopulation after exposure to a sub-lethal dose of IR[38, 39]. Our work uses *S. mediterranea* as an experimental model for assessing the resilience of IR-induced DNA damage and repair in adult tissues at the organismal level.

This study provides evidence that planarians can build tolerance for recurrent exposure to IR. We established a planarian colony expansion strategy to evaluate growth over successive rounds of sub-lethal radiation. We followed *S. mediterranea* growth and responses to IR exposure over four years (2014-2018). We also monitored animals for two more years after the last exposure to IR and did not register macroscopic tissue abnormalities. We discovered that while sub-lethal radiation led to phenotypic defects in a dose-dependent manner, higher levels of radiation exposure (17.5 Gy) resulted in increased expansion of colony size over time. Treating the colonies with a higher initial dose of sub-lethal radiation (i.e., 17.5Gy instead of 15 Gy or 10 Gy) helped them recover more effectively and better withstand population size in the first few weeks after multiple rounds of radiation. Additionally, we found that after five rounds of IR, the neoblast marker s*medpiwi-1* expression was not affected as in previous cycles of irradiation exposure. We also identified that levels of mitotic activity, DNA repair, and activation of HR through RAD51 nuclear translocation were enhanced over time. Thus, repetitive cycles of sub-lethal IR exposure led to developing RR in *S. mediterranea*.

## 2. Materials and Methods

### Planarian culture

The experiments described here used the clonal asexual strain CIW4 of *Schmidtea mediterranea*. Animals were maintained as previously described[40].

### Radiation exposure and colony expansion

Radiation exposure to planarian was administered using a Mark I Model 30 Cesium-137 irradiator. Animals were starved for at least one week before exposure to γ-radiation. To generate the initial colonies, ∼10-15 animals were exposed to varying sub-lethal doses of 10, 15, and 17.5 Gy previously characterized[23]. Colonies were allowed to recover to ∼20-30 animals for approximately 10-20 weeks post initial radiation exposure, depending on the recovery rate. Once recovered, colonies were split in half before radiation to generate the following: (1) a parental colony to maintain lineage tracking and generate an experimental control and (2) a daughter line with a subsequent exposure for experimental data collection. For subsequent exposures to generate daughter lineages, colonies were exposed to 15 Gy, as this is an intermediate level of sub-lethal irradiation that challenges neoblasts[23]. Still, most animals recover from it in a similar timeline (Figure 1C, D).

**Figure 1:**
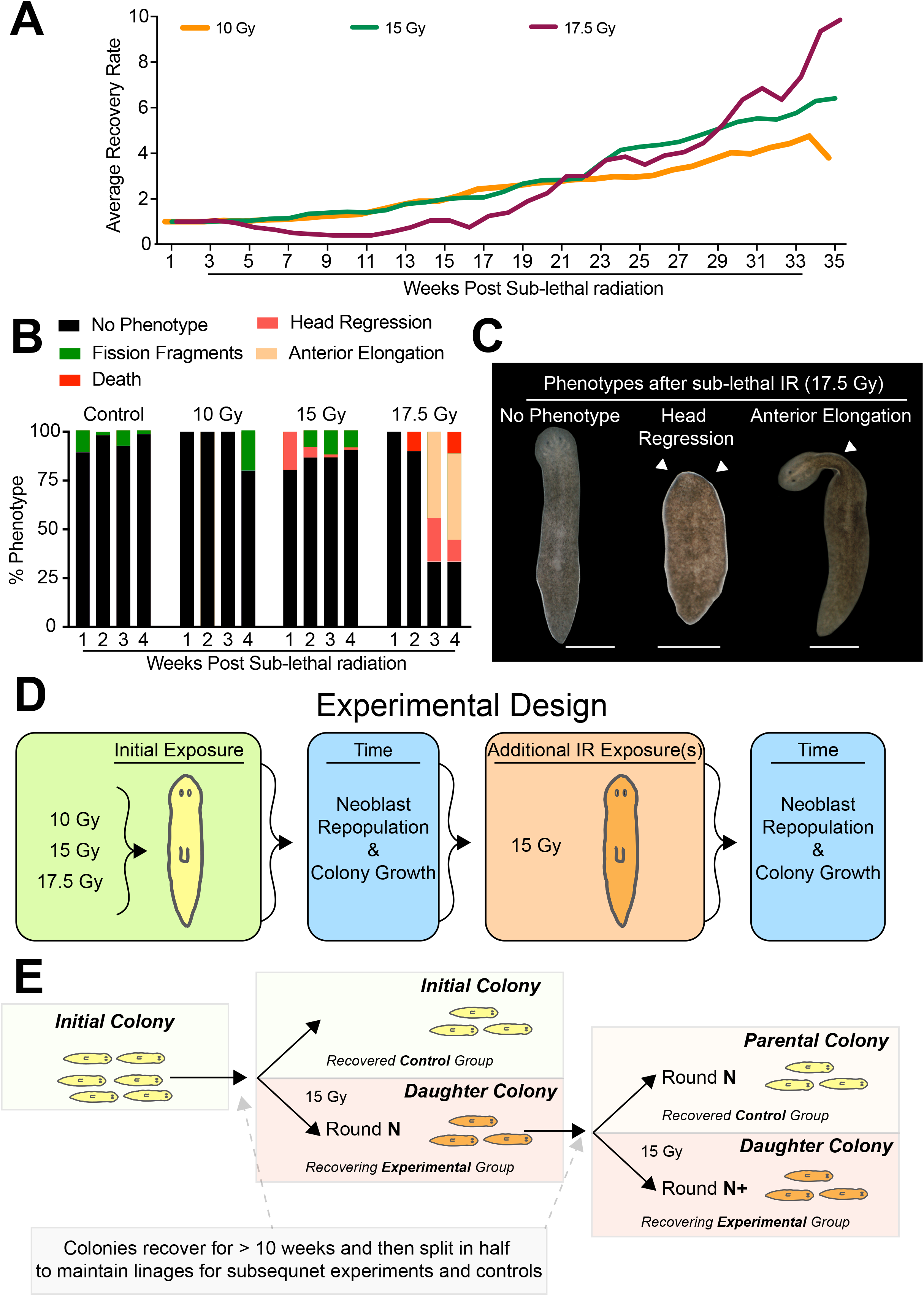
Planarian population recovery from sublethal ionizing radiation exposure facilitates long-term radio resistance experiments. **(A) The a**verage recovery rate of individual planarian colonies exposed to 10, 15, and 17.5 Gy. Averages represent 2-4 individual colonies consisting of 10-15 planarians per colony. Averages per colony can be identified in Figures S1A-C. **(B)** Phenotypes were recorded within the first four weeks post-radiation with 10, 15, and 17.5 Gy exposure relative to a non-irradiated control group. **(C)** Representative images of different phenotypes in the first four weeks after exposure to 17.5 Gy, scale bars = 200um. **(D, E)** Illustrations depicting the experimental strategy involving sub-lethal radiation and colony expansion. Note that the initial colonies are split in half after a > 10-week recovery period. This will maintain the initial colony and generate a daughter colony that will receive an additional round (round N) of 1.5K rads exposure. After an additional > 10-week recovery period, the daughter colony was split in half, forming a parental colony and a daughter colony (round N+), which received an additional exposure of 1.5K rads.

Furthermore, to generate colonies for molecular experiments, recovery periods were extended to account for colony split and replicates needed for each experimental assay and timepoint, needing tens to hundreds of animals per colony. In long-term experiments, animals recovered for > 24-52 weeks post-radiation. Colonies were split in half in all experiments, with the parental line undergoing long-term recovery and serving as the experimental non-irradiated control. The newly exposed daughter line was the experimental group to monitor short-term recovery.

### Colony growth rates and phenotypes

In short-term recovery experiments, colonies were monitored and cleaned weekly during the > 10-week recovery phase post-irradiation exposure. During this time, documentation of abnormal phenotypes and animal numbers was collected per colony. Strict numbering and colony tracking were followed over the years (Supplemental Figure 2-4). Thus enabling the monitoring of subsequent “lineages” originating from an initial parental colony. The calculation of colony growth rates is determined by the initial number of worms exposed to radiation. Thus, the current colony post-exposure number was divided by the input number of worms (i.e., number of worms/initial worm number). Note that if a subsequent exposure occurred during long-term colony recovery experiments, colony growth rates would be readjusted to account for the split in the colony. After the last IR exposure, we monitored colonies for two years to record any macroscopic abnormalities. Due to laboratory personnel rotation and the COVID-19 restrictions, the observations in most colonies were discontinued after 2020.

### Whole-mount and single-cell immunohistochemistry

Immunostainings were conducted as previously described for whole-mounts or dissociated single-cells[39, 41]. Briefly, 5-10 control and experimental radiation animals were fixed or dissociated before Carnoy’s fixation. Samples were blocked in 0.3%PBSTx+BSA for 4 hours, then incubated in primary antibody overnight, washed with 0.3%PBSTx, and incubated in secondary antibody. Primary antibody: α-Phospho-Histone H3 (Ser10) 1:250 (Millipore Cat# 05-817R), and α-RAD51 1:500 (Abcam, Cat# ab109107). Secondary antibodies: for α-H3P Alexa568 1:800 goat anti-rabbit (Invitrogen Cat# 11036). HRP-conjugated goat anti-rabbit antibody (Millipore, Cat# 12-348) was used at 1:500 for α-RAD51 with TSA-Alexa Fluor 488 (Thermo, Cat# A21370).

### Single-cell COMET assay

COMET assay under alkaline conditions allows DNA integrity analysis to be determined per individual planarian cells[39, 41, 42]. A pool of 5-10 worms was dissociated in CMF media and incubated at 37°C for a 2-hour recovery period. After recovery, cells were spun at 2500rpm for 2 minutes, resuspended in 140μL of 0.5% low melting point agarose, and spread on a 1% melting point agarose pre-coated frosted plus microscope slide. After the overnight lysis step, slides were neutralized in 0.4M Tris base pH 7.5 for 5 minutes at 4C. Slides were then subjected to 15 minutes of equilibration and 30 minutes of electrophoreses in prechilled 1x electrophoreses buffer (10N NaOH and 100mM EDTA, pH 13). Slides were then neutralized and fixed in 100% ethanol for 5 minutes. Once dry, slides were stained with a 1:10 ratio of SYBR gold (SYBR Gold; Invitrogen, Cat# S11494) into 1xTE buffer (10mM Tris-HCl, 1mM EDTA, pH 7.5) by placing 130μL of the solution onto the sample and covering with a cover slip (24×50mm). COMET tail lengths were determined using Nikon software.

### Whole-Mount *In-Situ* Hybridization

Expression levels of *smedpiwi-1* were determined by whole mount *in-situ* hybridization as previously described[43, 44].

### Image processing and analysis

Images obtained for phenotypes or immunohistochemistry of both cells and whole-mounts were processed using Nikon AZ-100 multi-zoom. Analysis of H3P or RAD51 positive foci was conducted by NIS Elements AR 3.2 or ImageJ software. Manuscript figures were prepared using Adobe Illustrator and Photoshop.

### Statistics and data representation

Data are expressed as mean ± standard error of the mean (SEM) or fold change ± SEM. For each experiment and timepoint, 6-8 animals from the control and treated groups were subjected to molecular experiments. Prism V10 software was used to generate graphs, and statistical significance was determined by Two-way ANOVA.

## 3. RESULTS

### 3.1. Planarian exposure to sub-lethal Ionizing radiation (IR) leads to dose-dependent phenotypic effects

We assessed organismal survival and recovery over time, following doses of sub-lethal radiation ranging from low to medium to high (i.e., 10, 15, and 17.5 Gy). We used four replicates for each sub-lethal dose group (10-15 animals each) to reduce the effects of variability in the response to IR. Upon IR exposure, animals within three colonies (one from the 15 Gy and two from the 17.5 Gy group) sustained irreversible damage and were eliminated. Thus, we monitored macroscopic tissue lesions and the recovery rate based on the increase in population size due to asexual reproduction through the fissioning of the remaining nine colonies. The results revealed that within the first four weeks post-irradiation (WPIR), the number of animals within each colony remained stable. Nonetheless, after five WPIR, the animals exposed to 10 and 15 Gy gradually expanded their colony. Conversely, planarians exposed to 17.5 Gy showed a reduction in the colonies’ population size for the next six weeks and began recovering after 15 WPIR. Independently of the initial dose of IR, all groups displayed similar levels of growth by 21 WPIR and thereafter (Figure 1A).

Exposure to sublethal doses of IR led to gross morphological defects. However, the severity of these defects (i.e., elongated pre-pharyngeal region, head regression, and death) appeared to be dose-dependent (Figure 1B, C). Animals exposed to 10 Gy appeared similar to control animals except that for one of the colonies at four WPIR, over 30% of the animals fissioned (Figure 1B). On the other hand, about 20% of the animals exposed to 15 Gy gradually lost anterior tissue (i.e., head regression) in the first WPIR. As the weeks progressed, a small fraction displayed a similar phenotype accompanied by animal fissioning. However, the group exposed to 17.5 Gy showed the most diverse range of defects that affected about 70% of the colony by the third WPIR (Figure 1B, C). These results confirm that tissue damage upon exposure to IR is dose-dependent. For this study, we will describe these nine colonies as the “initial colonies.”

Our results demonstrated that animals exposed to low and medium doses of sub-lethal radiation (15 Gy) could actively recover in weeks, which prompted us to examine the recovery rate after successive rounds of IR at a dose of 15 Gy (Figure 1D). Primarily, the initial colonies exposure to varying sub-lethal doses (10, 15, 17.5 Gy) were allowed to recover colony numbers for greater than 10 WPIR. After recovery, colonies were split in half; one was the parental colony, and the other was a daughter colony (Figure 1E). Here, one-half of the colony would receive an additional round of mild sub-lethal irradiation exposure (i.e.,15 Gy) to become a daughter colony, which would subsequently be used for data collection and lineage expansion. Furthermore, the other half of the colony would serve as a reference control (parental colony) with no additional IR exposures. The parental colony served as a node for lineage tracking and rapid expansion for future studies. Based on this experimental design, we expanded the initial nine colonies to over 47 colonies exposed to different rounds of radiation and monitored for over 3 to 4 years between 2014-2018 (Figures S2-4). These experiments were designed to evaluate the dynamics of the DNA repair and neoblast repopulation after successive rounds of radiation.

### 3.2. The initial dose of ionizing radiation exposure influences population dynamics and DNA integrity

To understand how multiple rounds of radiation impacted the long-term ability for population dynamics (asexual reproduction), colony size was recorded for more than 20 weeks following each round of 15 Gy exposure (Figure 2A). These results showed that the average colony size recovered at similar rates during the second round of 15 Gy exposure to what was observed after just one round of radiation, except for the initial colonies exposed to 17.5 Gy (Figure 1A, 2B). However, this recovery rate began changing during the third and fourth rounds and was reduced during a fifth round in both the 15 Gy and 17.5 Gy initial exposures (Figure 2B). Regardless of IR dosing, we observed an initial halt or slight decrease in growth, which began to increase between 6 and 10 WPIR (Figure 2B). These results suggested that population dynamics change in response to IR, with the initial dose influencing long-term colony expansion.

**Figure 2:**
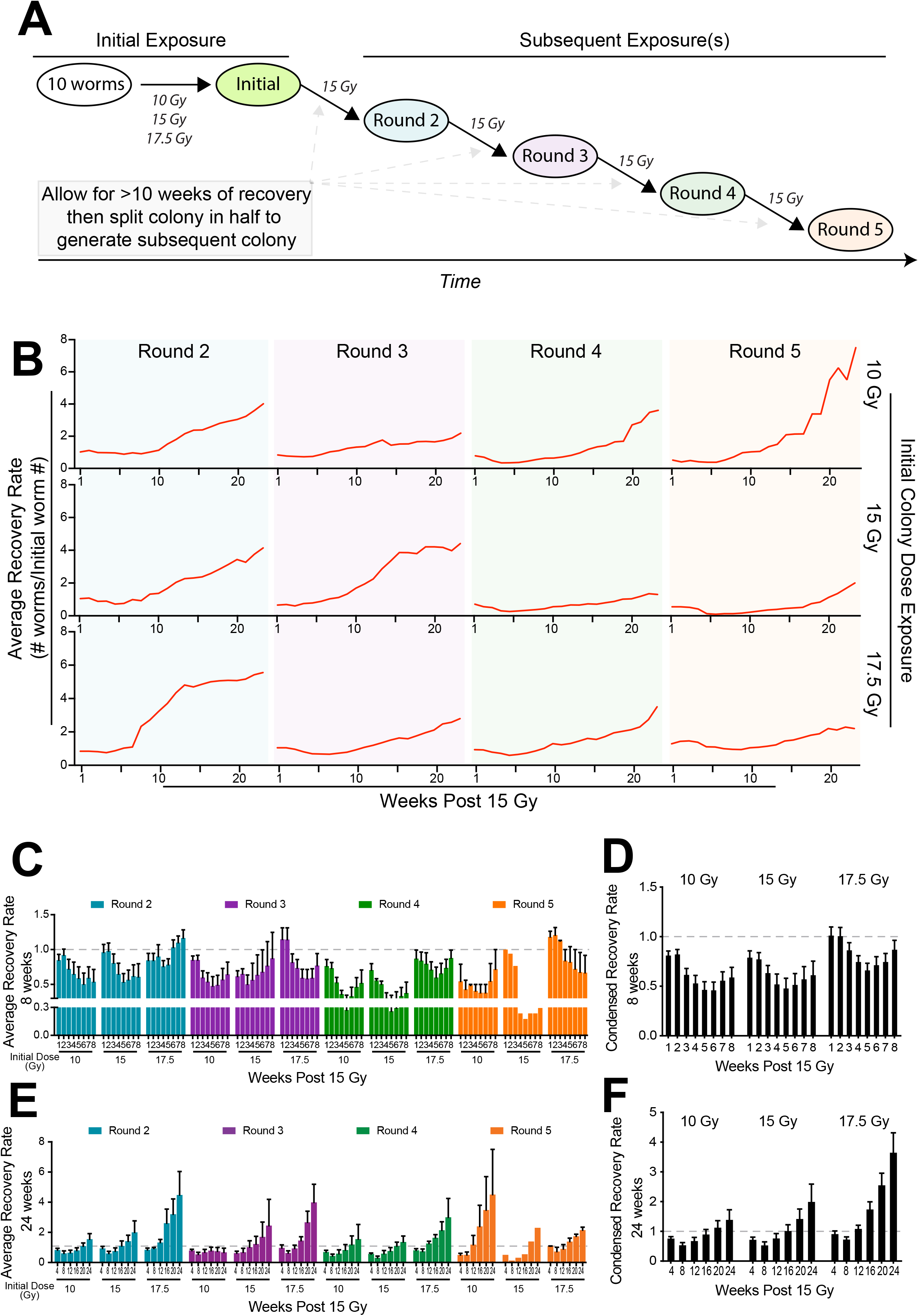
Planarian colony growth after recurrent exposure to sub-lethal ionizing radiation. **(A)** Schematic depicting the experimental strategy for colony expansion involving multiple rounds of sub-lethal radiation with 15 Gy. **(B)** Planarian colony growth over 20 weeks post-radiation. Data represents an average of all daughter colony recovery rates (e.g., N+ rounds) after subsequent rounds of exposure to 15 Gy. Average recovery rates were derived from Figures S5-7. **(C)** Average recovery rate during the first eight weeks for each round of ionizing radiation exposure. **(D)** The condensed average weekly recovery rate of all data points from data shown in C. **(E)** Average recovery rate during the first 24 weeks for each round of ionizing radiation exposure. **(F)** The condensed average weekly recovery rate of all data points from data shown in E. **(C-F)** Graphs represent mean ± SEM, and in most experiments, >100 animals were generated.

To better understand how these population dynamics were regulated, we compared technical replicates side by side to assess how radiation exposure influenced short-term (first 8 WPIR) and long-term (up to 24 WPIR) colony recovery. The short-term events within the first eight WPIRs suggested that the response to irradiation was not homogenous among the experimental groups but showed a similar trend among repeated rounds of exposure, which is evident by averaging the rounds of exposure for each experimental group (Figure 2C, D). We found that the initial response to irradiation (3-5 WPIR) led to a ∼ 30-40% reduction in colony size, which was immediately followed by a gradual recovery (Figure 2D). Additionally, planarians initially exposed to 17.5 Gy could withstand the first two WPIR better than the other two groups exposed to lower radiation levels (Figure 2C, D). We also found similar trends among rounds 2-4 across the experimental groups in the long-term colony recovery (Figure 2E). When averaging each experimental group across multiple rounds of exposure, we found that increasing the initial dose of radiation led to an increase in overall colony recovery (Figure 2F). These data suggest that despite the higher levels of tissue damage, the animals initially exposed to the higher dose of sub-lethal radiation could better withstand the initial response and long-term colony recovery.

Next, we assessed the overall colony recovery and the status of DNA integrity upon exposure to sub-lethal doses of IR. The COMET assay was used to determine DNA fragmentation, in which the length of the COMET tail is proportional to the degree of DNA fragmentation[42]. To generate a larger pool of animals per colony to allow for experimentation without depleting the colonies, we extended the recovery period to > 24 WPIR. By 24 weeks, animals have fully recovered and are indistinguishable based on the initial or the number of subsequent exposures (Figure S5). The fully recovered colonies (> 24 WPIR) served as controls (long-term) and were split in half to generate daughter colonies newly exposed to doses of 15 Gy. They were measured the following week (short-term) (Figure 3A). As expected, we found significant increases in DNA fragmentation in colonies recently exposed to irradiation (short-term) compared to colonies with 24 weeks to recover from irradiation exposure (long-term), suggesting that DNA damage was repaired over time. Within the same recovery lengths, levels of DNA fragmentation were similar regardless of the total number of exposures, except for the colonies receiving an initial exposure of 17.5 Gy (Figure 3 B-D). We found the (short-term) dynamics changed in colonies receiving 17.5 Gy; the third-round exposure colonies displayed slight increases in DNA fragmentation, while the fourth-round exposure colonies showed decreased levels after exposure (Figure 3 B-D). These findings suggest that the enhanced recovery rate in animals with an initial 17.5 Gy may be accompanied by increased DNA repair to restore DNA integrity.

**Figure 3:**
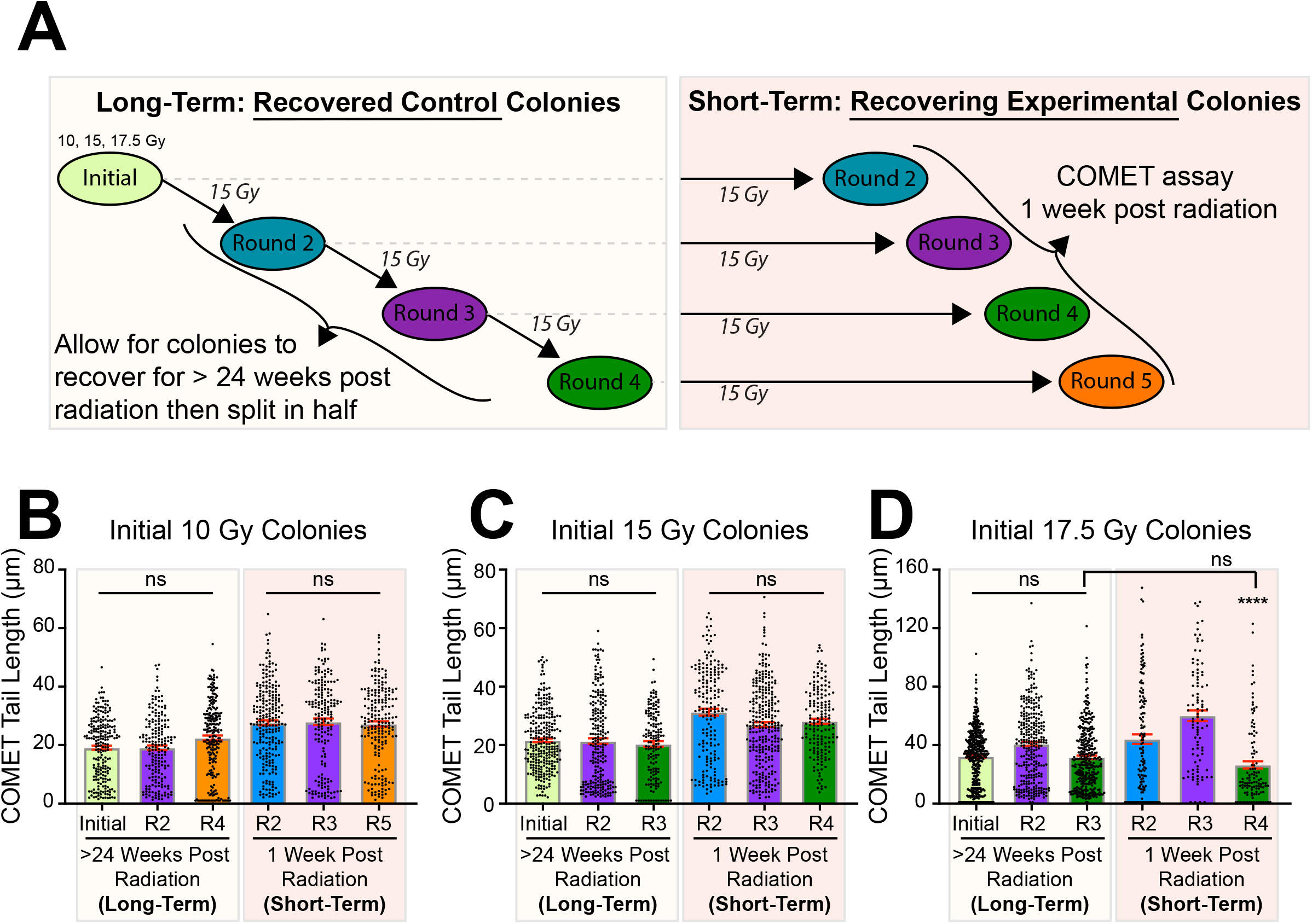
Planarians withstand the effects of radiation-induced DNA damage. **(A)** Schematic depicting the experimental strategy to evaluate DNA integrity with the COMET assay at short-term (one week) or long-term (24 weeks) post-exposure to three doses of sub-lethal radiation. The experimental colonies for the long-term experiments were split in half after each radiation round, with one group receiving subsequent N+ rounds of 1.5K rads exposure. **(B-D)** Data points indicate measurements of individual cell COMET tail lengths of initial exposure at 10, 15, and 17.5 Gy, respectively. For simplicity, each short- and long-term group was clustered and color-coded. In all cases or otherwise noted, radiation significantly increased tail length (**** < 0.0001) compared to the long-term recovered controls. COMET assay was conducted using five dissociated animals per time point, and each experiment per graph was generated and performed on different dates. *P*-value **** < 0.0001 and NS no significance were calculated by two-way ANOVA. Graphs represent mean ± SEM involving a preparation with five pooled animals.

### 3.3. The duration between multiple rounds of radiation exposure does not impact the short-term colony recovery

Based on our previous results, which show that the initial exposure of 17.5 Gy was associated with apparent protection in the colony size, long-term recovery rate, and enhanced DNA damage repair, we wanted to validate these findings in a larger cohort. Thus, to assess molecular events within the 17.5 Gy initial exposure colonies, we extended the recovery rate from 24 WPIR to 52 WPIR to obtain sufficient animal numbers per colony (i.e., greater than 150 animals per colony). To do this, we focused on colonies exposed to 2, 3, and 4 rounds with 15 Gy (Figure 4A). After 52 weeks, we split the colonies in half to generate mother and daughter colonies and subjected the daughter colonies to 15 Gy generating colonies with 3, 4, and 5 rounds of exposure.

**Figure 4:**
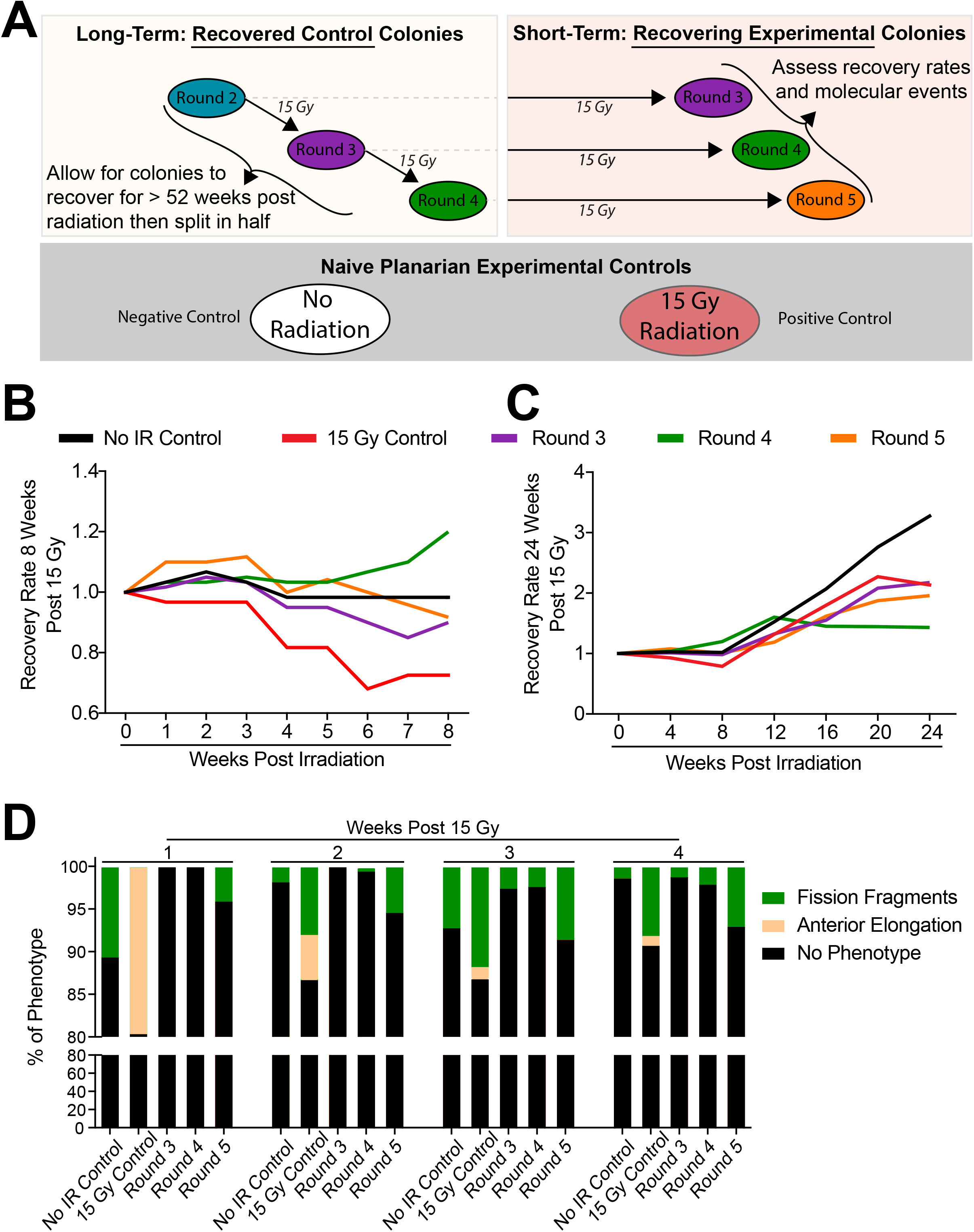
Increased rounds of ionizing radiation exposure result in accelerated colony recovery. **(A)** Schematic depicting the experimental strategy to generate planarian colonies that recovered 52 weeks after successive rounds of sub-lethal radiation. After recovery, colonies were split in half; one served as the control, and the other half served as the experimental radiation groups. Additionally, a positive and negative control composed of naïve planarians was used within this experiment to determine rates and depth of recovery. **(B-C)** Fold change in colony recovery rates over 8- and 24-weeks, respectively. Rates were juxtaposed against the negative and positive control groups. **(D)** Phenotypes were recorded within the first four weeks post-radiation with 15 Gy exposure relative to a non-irradiated control group.

Here, we assessed the short-term (first 8 WPIR) and long-term (extending to 24 WPIR) colony recovery dynamics. Furthermore, naïve planarian controls were implemented as a non-irradiated (negative) control and a 15 Gy exposure (positive) control. These controls served as baselines to compare known population recovery and repopulation dynamics of planarian cells post-radiation (i.e., mitosis, neoblast markers, and DNA damage dynamics) (Figure 4A). When monitoring colony growth, we identified that the colonies exposed to 15 Gy for over three rounds maintained a relatively stable recovery rate compared to the naïve 15 Gy control in the early WPIR (Figure 4B). Short-term recovery increased during the first three WPIR in animals exposed to five rounds of radiation and 3-8 WPIR in animals exposed to four rounds. The high recovery rate during the first 12 WPIR in animals exposed to five rounds of radiation was similar to animals exposed to 15 Gy during the first two WPIR (Figure 2C, D, 4C). However, these recovery rates dropped for the long-term (> 12 WPIR) recovery. After 12 WPIR, independent from the amount of exposure, the recovery rate was decreased compared to control, non-irradiated animals (Figure 4C). These results suggest that regardless of the length of recovery, 24 WPIR vs. 52 WPIR, the short-term recovery rate is maintained longer in animals receiving increased radiation exposure. However, long-term recovery (>24 WPIR) decreased; this was most evident in animals receiving an initial 17.5 Gy after four rounds of radiation (Figure 2D, E 4C). We determined that this increased early recovery rate directly resulted from increased fission fragments (Figure 4D).

### 3.4. Neoblasts develop resilience to withstand recurrent rounds of ionizing radiation exposure

To gain a deeper insight into the cellular and molecular process allowing planarians to recover following multiple rounds of radiation, we analyzed neoblast gene expression, mitotic activity, and dynamics of RAD51, a protein involved in DSB damage repair. The neoblasts are the only cells with the capacity to divide and possess the ability to differentiate into any planarian cell type[23-27]. In response to sub-lethal radiation exposure, neoblasts are temporarily depleted, followed by a gradual repopulation[32, 37]. Using whole-mount *in* situ hybridization with the neoblast maker *smedwi-1*, we could visualize the spatial neoblast recovery in animals 1, 2, or 3 WPIR after repeated rounds of exposure (Figure 5A). Qualitatively, these results showed a reduction of *smedwi-1* expression even after two WPIRs. However, after three WPIR, the expression of *smedwi-1* resembled that of the unirradiated animals. This impact was most dramatic in animals exposed to three and four rounds of radiation.

**Figure 5:**
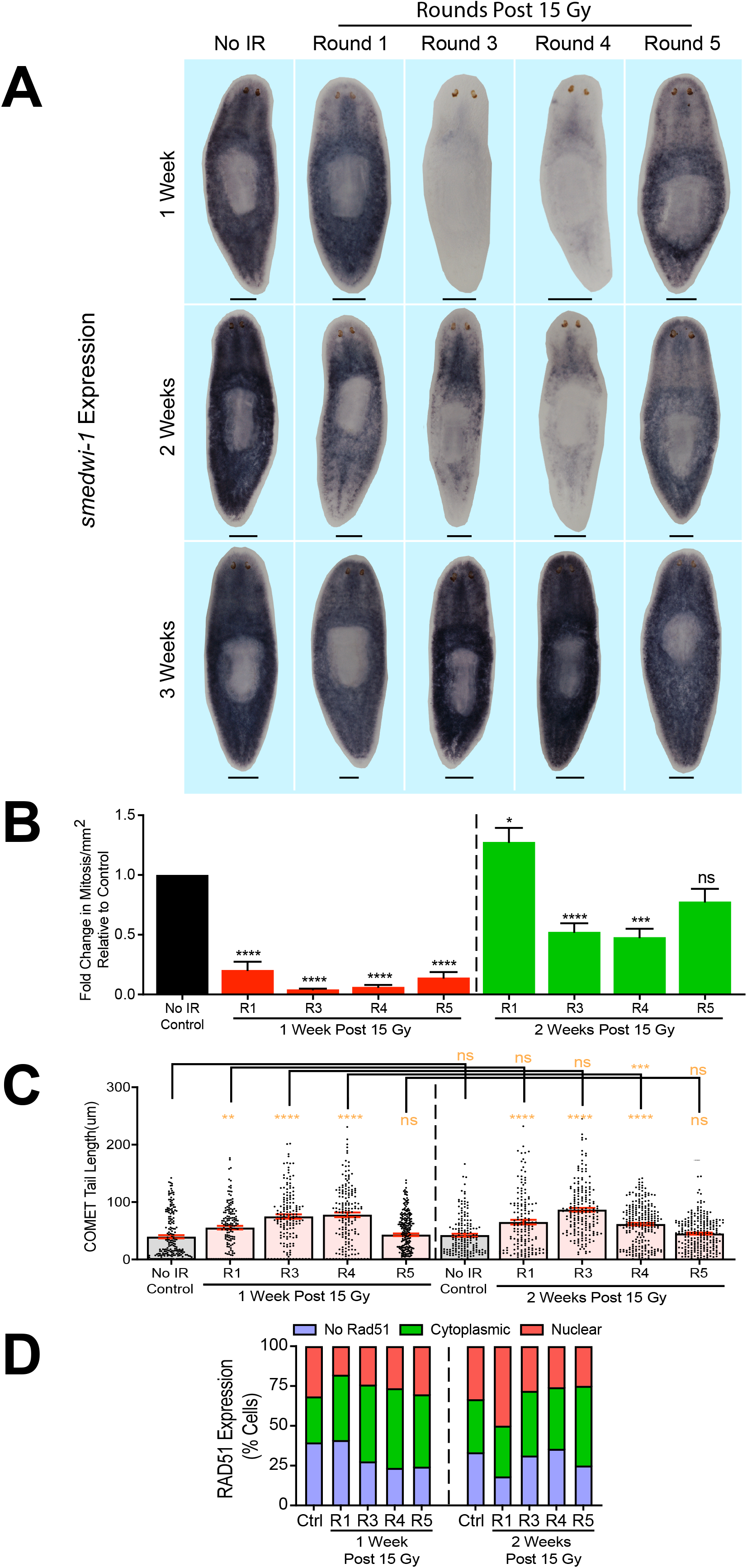
Neoblasts develop resilience to repeated rounds of sub-lethal ionizing radiation. **(A)** Representative whole-mount *in-situ* hybridization images with the neoblast marker *smedwi-1* (purple signal) after exposure with 15 Gy. Eight samples were fixed weekly for three weeks. Scale bars= 200μm. **(B)** Mitoses levels were obtained by whole-mount immunostaining with the phospho-histone H3 (Ser10) (H3P) antibody at one- and two weeks post 15 Gy exposure. Immunostaining was conducted on ten animals per time point and condition. **(C)** DNA integrity was measured with the COMET assay. Data points indicate individual cell COMET tail lengths after one- or two-week radiation exposure with 15 Gy. **(D)** Cytoplasmic or nuclear RAD51 expression on dissociated cells at one- and two weeks post-radiation 15 Gy. The signal location was recorded based on its location in the cytoplasm or nucleus in each cell. **(C-D)** Single cells were derived from five dissociated animals per time point and condition. **(B-C)** Graphs represent mean ± s.e.m. *P*-values * <0.05, ** < 0.001, *** < 0.0005, and **** < 0.0001 were calculated by two-way ANOVA.

Interestingly, the reduction in *smedwi-1* expression seemed less dramatic in animals with five rounds of IR exposure (Figure 5A). These results suggest that the effects on *smedwi-1* expression are less pronounced after five rounds of IR. Next, we wanted to evaluate if the dynamics of neoblast division display any change after several rounds of radiation. Thus, we monitored mitotic activity at one and two WPIR, coinciding with the reduced *smedwi-1* expression (Figure 5A-B). As expected, the rate of cellular division was significantly diminished by one WPIR independently of the rounds of IR exposure (Figure 5B). Nonetheless, we observed a striking rebound in cell division during the second WPIR. Animals with one round of IR exposure had higher levels of cell division than the unirradiated animals. The groups receiving three and four rounds showed an increase in cell division, half of the control samples. Interestingly, the group with five rounds of radiation displayed similar levels to unirradiated animals by the second WPIR, suggesting these animals could reestablish cell division faster than the other experimental groups (Figure 5B).

Next, we wanted to evaluate changes in the levels of DNA damage over repeated exposure to IR. We found similar trends in one and two WPIR animals, in which DNA damage was significantly increased between one and four rounds of IR exposure. However, animals subjected to five rounds of irradiation consistently showed similar levels of DNA damage as the non-irradiated control (Figure 5C). We noticed that COMET tail length for animals with four rounds of irradiation significantly decreased by two WPIR, suggesting that DNA integrity recovers faster after the second-week irradiation cycle in this group.

Nuclear translocation of RAD51 is necessary to reestablish DNA integrity upon IR exposure[39]. We identified that at one WPIR, RAD51 expression was elevated in the cytoplasmic fraction across all radiation-exposed groups, and the nuclear fraction, which was initially decreased, showed stepwise increases upon increased rounds of radiation exposure (Figure 5D). Interestingly, our positive control (Round 1) did not show an increased nuclear RAD51 expression relative to the non-radiated control. However, by two WPIR, the positive control group increased expression of total RAD51 and was elevated within the cell nucleus. At this time, cells derived from colonies with more than three rounds of irradiation seemed to reestablish RAD51 expression distribution comparable to the control group (Figure 5D). Thus, a more fluid nuclear-cytoplasmic dynamic was inferred with increased irradiation rounds. These data suggest that after repeated rounds of radiation exposure, neoblasts have either resistance to ionizing radiation-induced DNA damage and/or acquired the ability to implement a more effective DNA damage repair. We inferred DNA functional fitness based on the long-term recovery dynamics of body size (surface area), cellular proliferation rates, and regenerative capacity (Figure S5). We found that regardless of initial exposure doses or repeated rounds of exposure, their long-term recovery dynamics were similar (Figure S8). This implies that the dynamics of neoblasts exposed to radiation can adapt, allowing animals to maintain colony growth despite repeated DNA insults.

## 4. Discussion

Our results show that planarians can adapt to overcome the detrimental effect of recurrent IR exposure. We found comparable DNA damage levels between unirradiated controls and long-term recovery animals exposed to multiple rounds of sub-lethal IR. This finding supports the possibility of strengthening cellular and molecular strategies to protect and repair DNA damage when subjected to repeated insults with IR. We also identified that animals exposed to higher levels of sub-lethal irradiation (17.5 Gy) during the first exposure were better at withstanding the initial response and long-term recovery to irradiation. This was more prominent after multiple rounds of irradiation exposure, which we propose was partly due to a more resilient response to repair DNA.

The planarian-indeterminate life cycle enables long-term studies investigating DNA repair resilience to repeated insults. Planarians can live for years by continuously undergoing tissue renewal driven by neoblasts[29, 30, 45]. Thus, the extended life cycle may entail efficient DNA repair mechanisms to sustain the structural DNA integrity required to support cellular turnover fueled by continuous neoblast division. Following exposure to IR, lesions are created in the DNA, and the DSBs are the most significant[4]. The planarian genome exhibits evolutionarily conserved pathways to repair DSBs[38]. Functional analyses demonstrated that components associated with HR through RAD51 are the predominant pathway involved in DSB repair in planarians[37-39, 41]. We observed no gross macroscopic changes after two years of the last IR exposure, and together with the sustained capacity to properly grow, regulate mitosis levels, and regenerate tissues upon amputation, we inferred that the long-term functionality of neoblasts was maintained in planarians exposed to five rounds of sub-lethal IR (Figure S5). Due to the sizeable pluripotent population of stem cells and conservation of DNA repair components, planarian flatworms are an attractive model for studies on RR development and DNA repair in the context of continuous cellular turnover in the adult body.

Planarians can withstand higher radiation doses than most other mammals[46, 47]. In response to sub-lethal doses (i.e., below 17.5 Gy), there is a transient loss of neoblasts within 24 hours, followed by a gradual repopulation across the body[23, 48]. It has been proposed that EGF signaling and DNA repair components, such as ATM, RAD51, Ku70, and RAD54B, mediate the neoblast repopulation[39, 48]. Our findings suggest that after several rounds of sublethal radiation, DNA damage, and the RAD51 nuclear translocation are comparable to those of the unirradiated control group. This is accompanied by equivalent levels of neoblast division between the control and experimental groups, implying that the effects of IR are minimized over time. Future studies will address whether DNA protective mechanisms, as shown in tardigrades, contribute to the resistance to DNA damage upon IR[12]. Furthermore, testing whether planarian-specific molecules are involved in RR would be interesting. Recent work has shown that transferring tardigrade-specific proteins, such as Tdar, can protect human cells against IR[20]. These findings suggest evolutionary conservation and potential translational applications geared toward clinical intervention with radiotherapy.

The molecular details driving radiotolerance in planarians need to be elucidated. Nonetheless, given that RAD51 nuclear translocation is enhanced after IR, we propose that at least HR signaling pathways may be involved in the process of radiotolerance in planarians. Future studies may include transcriptome analysis on animals exposed to multiple rounds of IR to dissect the mechanisms of RR in planarians. Recent work shows that short stimulation with direct current stimulation could enhance DNA damage repair and promote stem cell repopulation in planarians lethally irradiated[37]. This suggests that DNA repair mechanisms could be enhanced and directed toward stem cell populations necessary to reestablish cellular turnover in adult tissues. In addition, previous transcriptomic analysis uncovered a subset of genes upregulated upon lethal irradiation and suggested possible signaling pathways involved in sensing DNA damage and the signal transduction triggered by DSBs in planarians[33, 49, 50]. It is tempting to assess the hypothesis that genes overexpressed after lethal irradiation may also contribute to the planarian adaptation to effective DNA repair after several cycles of sub-lethal irradiation.

Enhanced DNA damage repair under extreme stress is not specific to only IR exposure. The Blind mole rat, *Spalax ehrenbergi, and the naked mole rat, Heterocephalus glaber*, can survive under extreme conditions such as low-oxygen environments[51, 52]. Typically, these conditions would lead to reactive oxygen species (ROS)-mediated DNA damage and genomic instability; however, enhanced DNA damage repair abilities have allowed for genotoxic stress resistance[52, 53]. As a result, both species have a rare occurrence of spontaneous cancer, enabling them to be long-lived organisms compared to other rodents[54-57]. In fact, upon treatment with carcinogens DMBA/TPA and 3MCA, *S. ehrenbergi* either did not develop tumors or had a low frequency of tumors over extended periods[56, 57]. Planarians also exhibit evolutionarily conserved ROS-responsive systems and show a low incidence of spontaneous tumors under rearing lab conditions[58-61]. Nonetheless, cancer-like phenotypes could be elicited in planarians by disturbing the functions of tumor suppressor genes[62]. Despite these susceptibilities, *S. mediterranea* has a robust strategy for preserving genomic integrity, allowing them to be long-lived organisms[38, 45]. It is possible that planarians already have an enhanced DNA damage repair system compared to other organisms. These mechanisms can adapt to even more efficient responses upon extreme stresses such as repeated low-dose radiation exposure or stimulation with electric currents[37]. Future analysis of animals with enhanced DNA repair mechanisms may help understand molecular mechanisms of acquired radio tolerance in adult organisms.

## 5. Concluding remarks

The results presented here offer unique opportunities to understand the development of radiotolerance. Our findings enable the possibility of designing population studies to select specific attributes, such as enhanced stem cell recovery from radiation. It will be interesting to probe for DNA repair fidelity and mutation rates after repeated exposure to IR. We have used animal growth and regeneration as readouts, but DNA sequencing may provide complementary information on the accuracy of planarian radiotolerance. Likewise, repopulation studies on planarians with cancer-like phenotypes [62] may help understand cancer stem cell dynamics to chemotherapy/radiotherapy in the complexity of the adult body. Studying the pathways cancer stem cells use to acquire resistance to chemotherapy/irradiation in mammals is challenging[63]. Therefore, we propose using planarians to uncover potential new avenues of radiation resistance that can otherwise be hidden in the multitude of mutations in mammals; further studies utilizing planarians and their neoblasts present fresh opportunities.

## 6. Acknowledgments

We thank members of the Oviedo lab for insightful discussions and comments on the manuscript. We are grateful to an extensive number of undergraduate students at UC Merced who assisted with the maintenance of planarian colonies over the years of these studies.

## 8. Author contributions

P.G.B., B.Z., E.M., N.M.F., P.K., S.R., E.P., E.M., and N.J.O. conceived, designed, analyzed data, data curation, maintained planarian colonies, and executed the experiments. P.G.B., B.Z., and N.J.O. drafted the original manuscript; all authors read, commented, and approved the final version. N.J.O. supervised and secured funding and resources for the execution of the project.

## 11. Supplemental Information

**Supplemental Figure 1: Planarian growth rates after exposure to sublethal radiation. (A-C)** The growth rate for each planarian colony exposed to a sub-lethal dose of ionizing radiation was recorded for sixty weeks. The average for all colonies was calculated and juxtaposed against the individual growth rate per line.

**Supplemental Figures 2-4: Recovery rates for planarian colony lineages exposed to recurrent sub-lethal ionizing radiation**. The graphs show a fold change in recovery rates based on the increase in population number over the initial group of animals that each group started with. Sub-lethal dose exposure (i.e., 10, 15, and 17.5 Gy) started with at least two groups (10-30 animals each). To illustrate the overall strategy, all groups for each IR dose are shown, but in each case, only one or two groups were followed for the total length of the study. Each “lineage” colony was named based on the IR dose and the line number that changed as the colony was split upon successive rounds of IR. Each group was linked to the initial colony and tracked over 130-150 weeks. The ionizing radiation exposures are indicated with arrows and their respective dose exposure.

**Supplemental Figure 5: Long-term recovery does not alter planarian homeostasis and regeneration**. Recovery measurements after more than 24 weeks post-radiation **(A)** Surface area **(B)** Levels of cell division determined by using the phospho-histone H3 (Ser10) (H3P) antibody **(C)** Surface area measurements of blastema size seven days post-amputation for animals initially exposed to 17.5 Gy. Animals were amputated into three fragments (i.e., head, trunk, and tail) and quantified as anterior or posterior blastemas. **(A-C)** Graphs represent mean ± s.e.m. *No significance (NS)* for all data points was calculated by two-way ANOVA. Five animals were used per timepoint and replicated once in all cases. R2-R5 denotes the rounds of irradiation (color-coded), and the numbers preceding the parenthesis indicate the labels assigned to each group under the lineage.

