## Supplementary figures and images for "Planarians Develop Radiotolerance to Recurrent Ionizing Radiation Exposure"

### Supp Figures

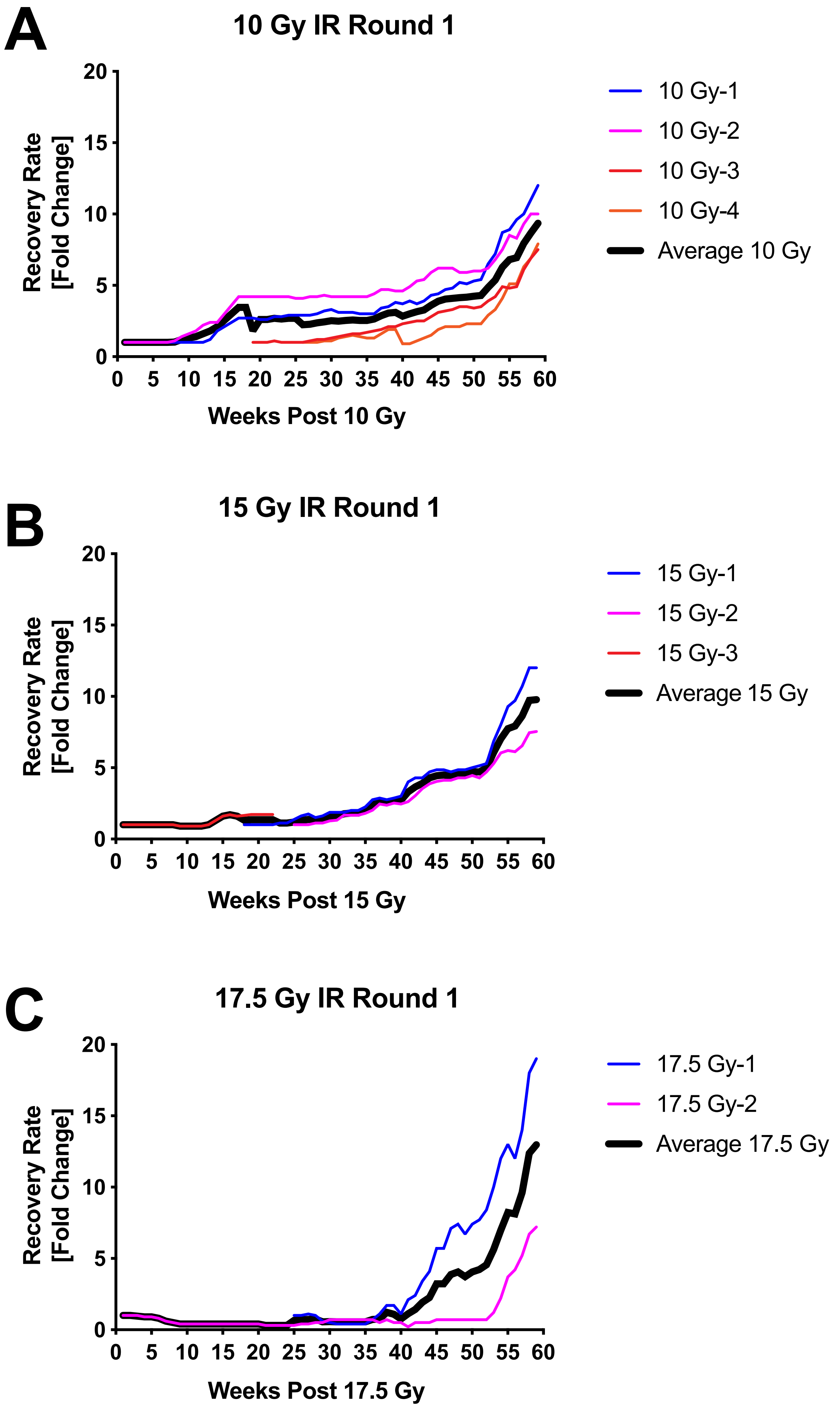

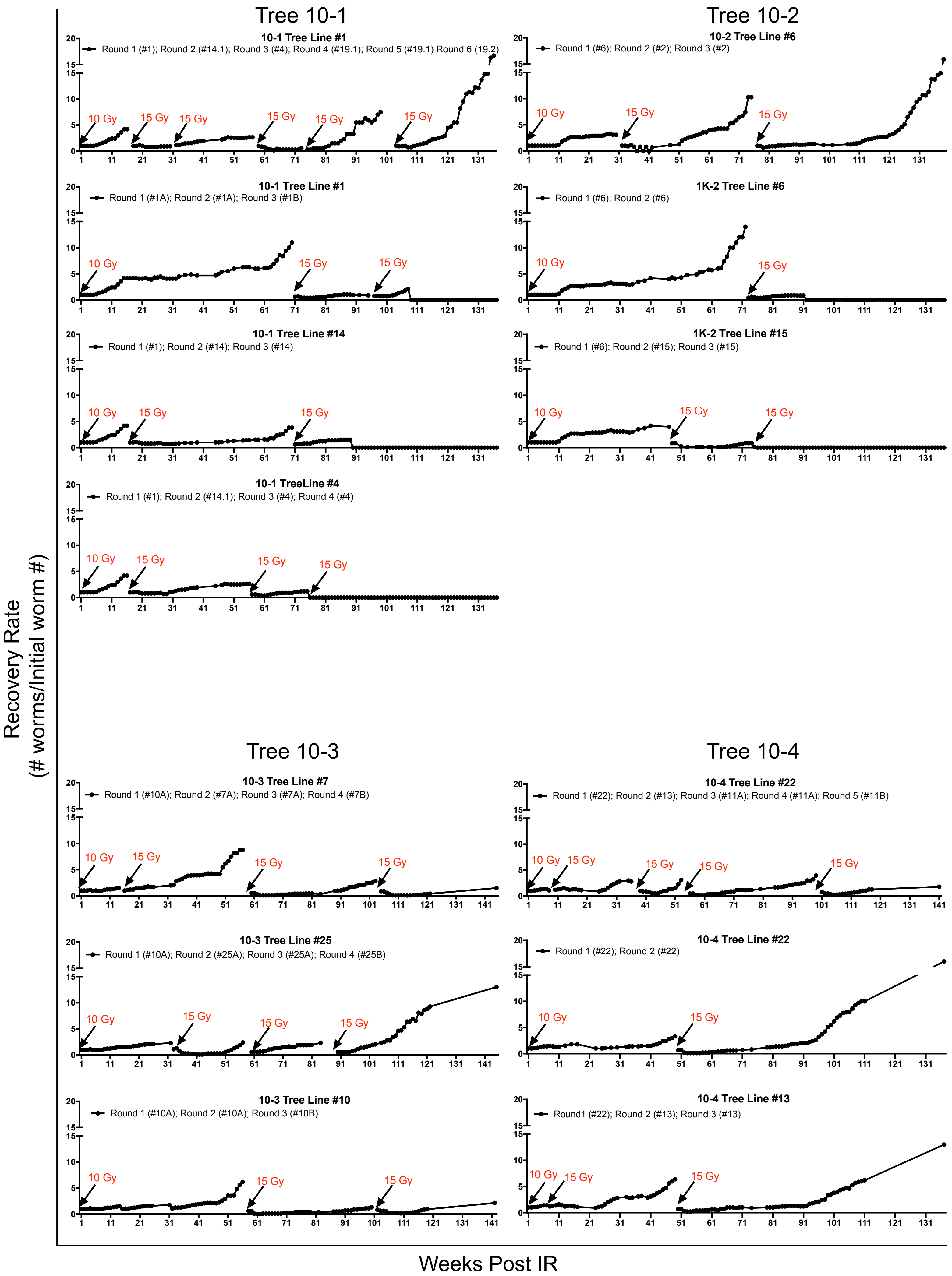

*Barghouth et al. Supplemental Figure 3*

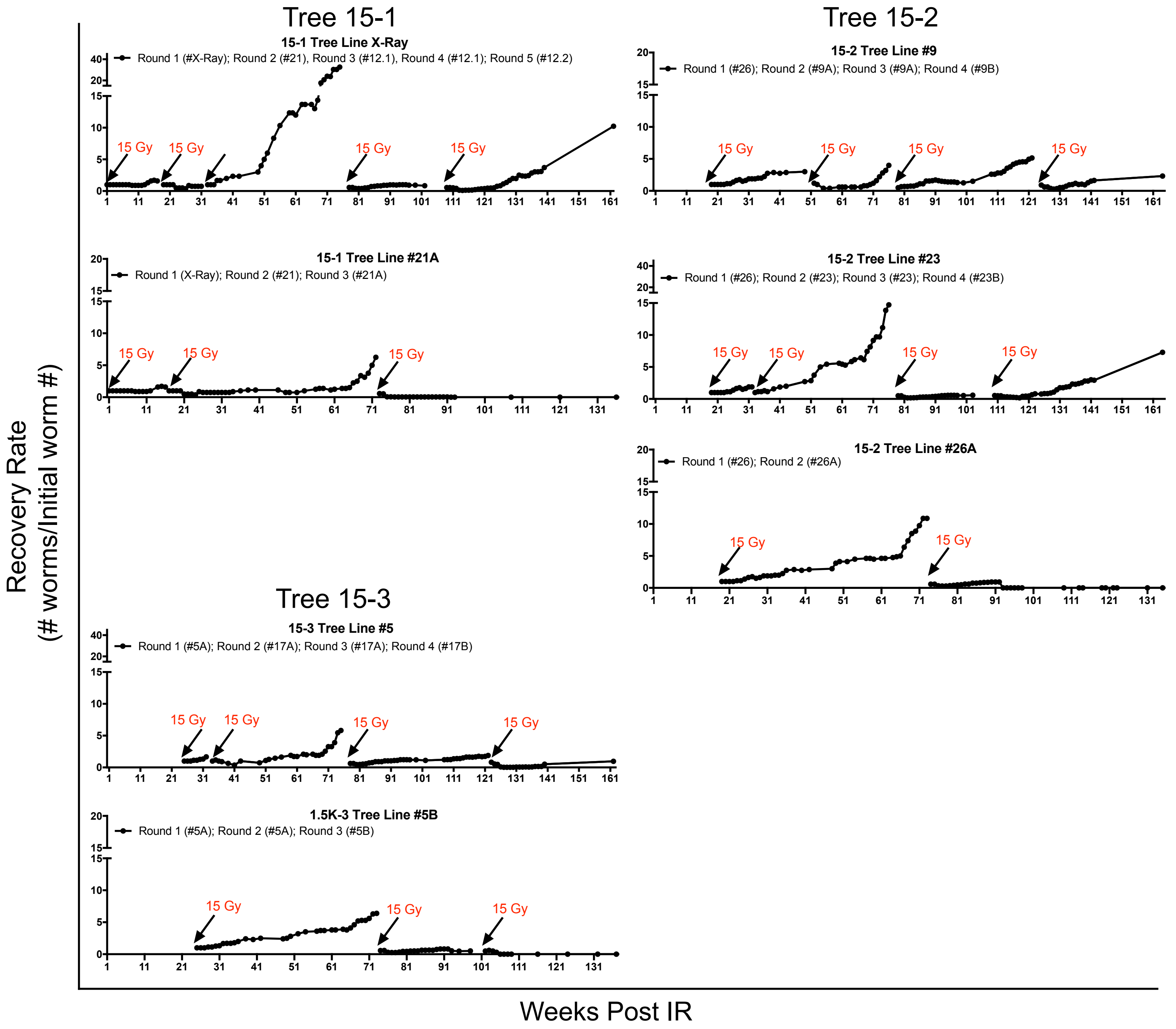

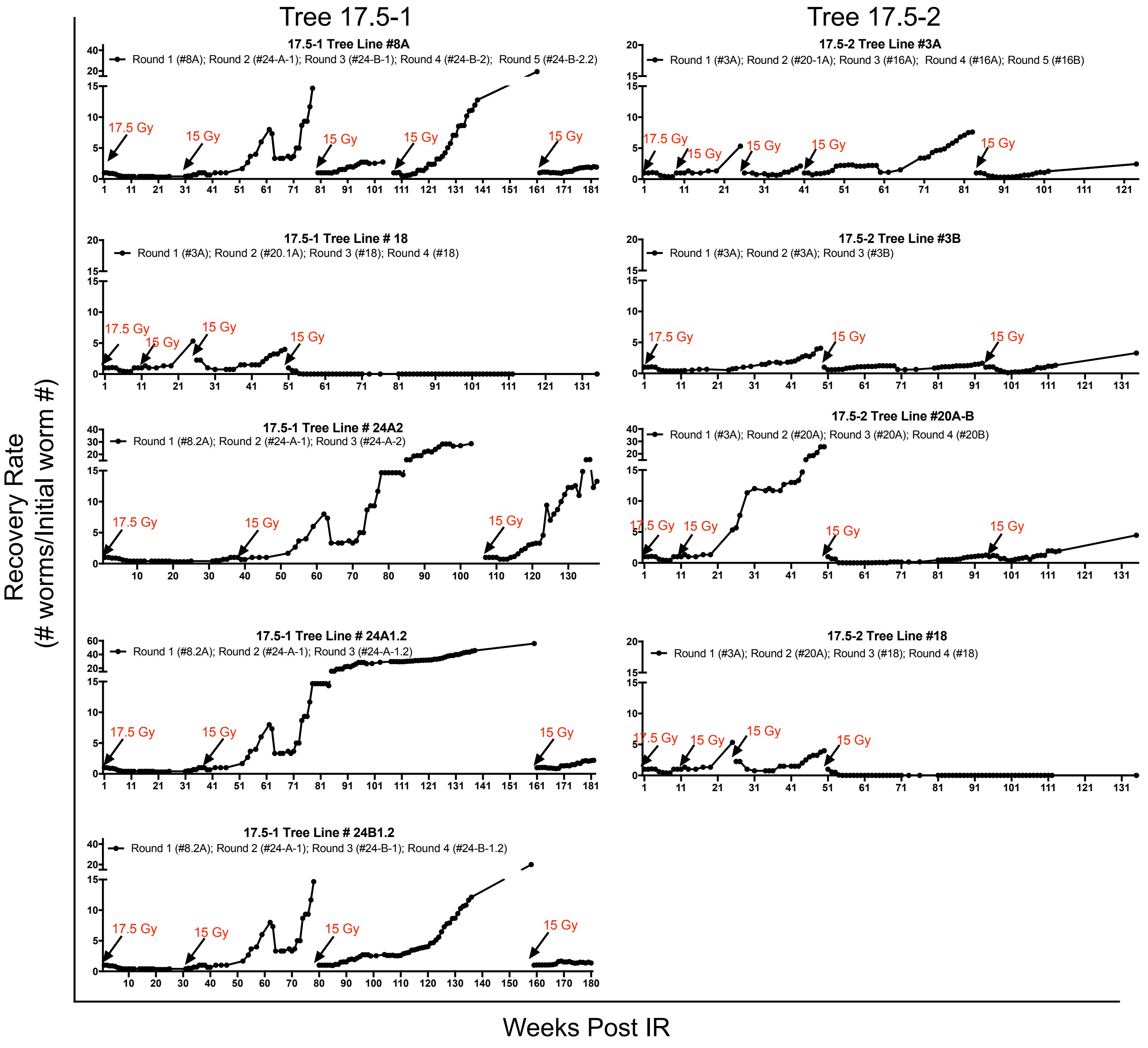

#### Long Term Recovery Dynamics (> 24 Weeks)

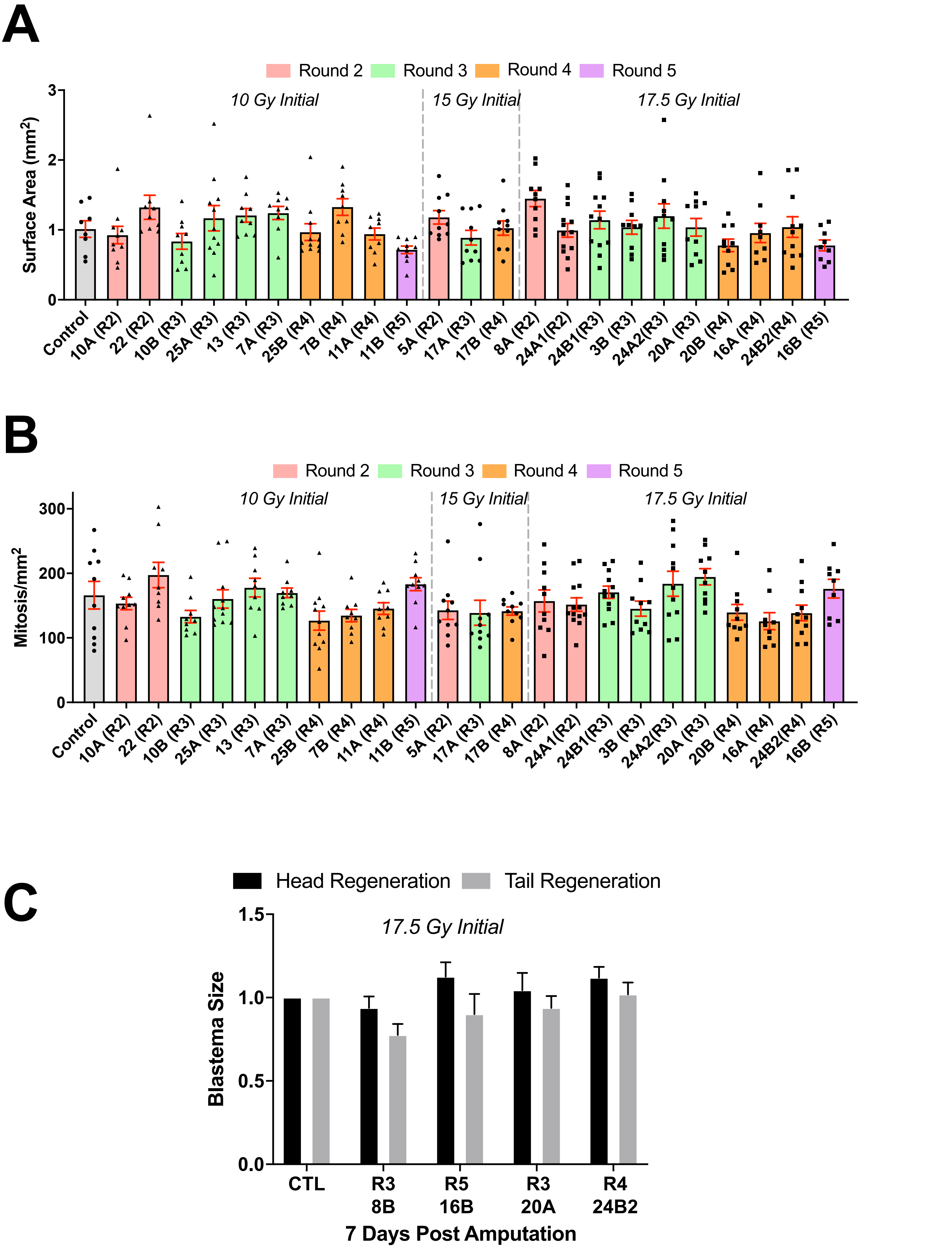
